# Fasting prevents hypoxia-induced defects of proteostasis in *C. elegans*

**DOI:** 10.1101/278671

**Authors:** Nicole N Iranon, Bailey E Jochim, Dana L Miller

## Abstract

Low oxygen conditions (hypoxia) can impair essential physiological processes and cause cellular damage and death. We have shown that specific hypoxic conditions disrupt protein homeostasis in *C. elegans*, leading to protein aggregation and proteotoxicity. Here, we show that nutritional cues regulate this effect of hypoxia on proteostasis. Animals fasted prior to hypoxic exposure develop dramatically fewer protein aggregates compared to their fed counterparts, indicating that the effect of hypoxia is abrogated. Fasting is effective at protecting against hypoxia-induced proteostasis defects in multiple developmental stages, tissues, and in different models of misfolded or aggregation prone proteins. Our data also demonstrate that the effect of fasting is induced and reversed quite rapidly, suggesting that the nutritional environment experienced at the onset of hypoxia dictates at least some aspects of the physiological response to hypoxia. We further demonstrate that the insulin/IGF-like signaling pathway plays a role in mediating the protective effects of fasting in hypoxia. Animals with mutations in *daf-2*, the *C. elegans* insulin-like receptor, display wild-type levels of hypoxia-induced protein aggregation upon exposure to hypoxia when fed, but are not protected by fasting. However, we found that DAF-2 acts independently of the FOXO transcription factor, DAF-16, to mediate the protective effects of fasting. These results suggest a non-canonical role for the insulin/IGF-like signaling pathway in coordinating the effects of hypoxia and nutritional state on proteostasis.

**Author Summary:** When blood flow to various parts of the body becomes restricted, those tissues suffer from a lack of oxygen, a condition called hypoxia. Hypoxia can cause cellular damage and death, such as is observed as a result of stroke and cardiovascular disease. We have found that in the model organism *C. elegans* (a roundworm) specific concentrations of hypoxia cause aggregation of polyglutamine proteins – the same kind of proteins that are found in an aggregated state in the neurodegenerative disorder Huntington’s disease. Here, we show that that worms can be protected from hypoxia-induced protein aggregation if they are fasted (removed from their food source) prior to experiencing hypoxia. Furthermore, we show that the insulin receptor is required for this protection. The insulin receptor is responsible for detecting insulin, a hormone that is released after feeding. Worms with a nonfunctional version of the insulin receptor displayed hypoxia-induced protein aggregation despite being fasted before the hypoxic exposure. Our results highlight a new role for the insulin signaling pathway in coordinating the effects of both hypoxia and nutritional state on protein aggregation.

## Introduction

In order to survive in changing conditions, organisms need to successfully integrate a number of environmental signals and respond appropriately in order to maintain homeostasis. Aerobic heterotrophs must meet their requirements for food and oxygen by taking in these resources from the environment. An inadequate response to low levels of oxygen (hypoxia) can lead to cellular damage or death, an unsurprising outcome given oxygen’s central role in cellular metabolism. Like hypoxia, food deprivation presents an obstacle to homeostasis by impinging on cellular metabolism and disturbing anabolic pathways. However, in many cases food restriction can have beneficial effects, such as extending lifespan and delaying the onset of neurodegenerative diseases and their associated pathologies [1]. In a mouse model of Alzheimer’s disease, 12 weeks of caloric restriction reduces Aβ plaque burden [2], and mice expressing human mutant huntingtin maintained on an alternate-day-feeding diet have reduced brain atrophy and decreased huntingtin aggregate formation [3]. Similarly, depriving *C. elegans* of their bacterial food source reduces damage associated with expressing polyglutamine proteins [4].

The protective effect of fasting is not limited to symptoms of neurodegeneration – there are many studies that show fasting can protect against damage associated with hypoxia in mammals. For example, mice on an alternate-day feeding regimen have higher survival rates after myocardial ischemia induced via coronary occlusion [5]. Similar results have been obtained with ischemic damage to the liver. Mice on a calorically restricted diet have reduced infarct damage compared to ad-libitum fed controls [6], and mice that have been fasted for 3 days display reduced hepatocellular apoptosis and damage [7]. Calorie restriction can also improve outcomes after cerebral ischemic injury by protecting cortical and striatal neurons [8], and reducing neurological deficits and infarct volume [9]. These observations suggest that understanding the mechanistic basis underlying the protective effects of fasting in hypoxia could provide novel insight into therapeutic strategies to treat pathological conditions associated with ischemia and reperfusion injury.

We have previously shown that in *C. elegans* the cellular response to specific hypoxic conditions involves a disruption of proteostasis – the coordination of protein synthesis, folding, degradation, and quality control required to maintain a functional proteome [10]. Here we show that fasting prevents the hypoxia-induced disruption of proteostasis. Our data indicate that the nutritional context of an animal at the onset of hypoxia has the power to alter hypoxia’s effect on proteostasis and that the insulin-like signaling (IIS) pathway plays a role in fasting’s ability to protect against proteostasis decline independently of the canonical downstream transcription factor DAF-16/FOXO.

## Results

In order to investigate the effect of nutritional status on proteostasis in hypoxia, we first used transgenic *C. elegans* that express yellow fluorescent protein (YFP) fused to a polyglutamine tract in the body wall muscles [11]. We refer to these animals as QX∷YFP, where X refers to the number of glutamine residues fused to YFP, such that Q35∷YFP animals express YFP with 35 glutamine residues. In these animals, the number of YFP foci, which correspond to large protein aggregates, can be used as an *in vivo* measure of cellular proteostasis [12].

Exposing animals to 0.1% oxygen for 24 hours while fed resulted in an increase in the number of YFP foci (Fig. 1B-1D), consistent with a decrease in proteostasis as has been demonstrated previously [10]. However, we found that the number of YFP foci that formed in hypoxia was dramatically reduced if the animals were removed from food for six hours before the hypoxic exposure and remained off of food for the duration of hypoxia (Fig. 1A). Hypoxia-induced protein aggregation (HIPA) was prevented by fasting in fourth-stage larvae (L4) Q35∷YFP animals (Fig. 1C) as well as in first-stage larvae (L1) Q40∷YFP (Fig. 1D). We conclude that fasting prevents HIPA and that this effect persists across development.

**Figure 1.**
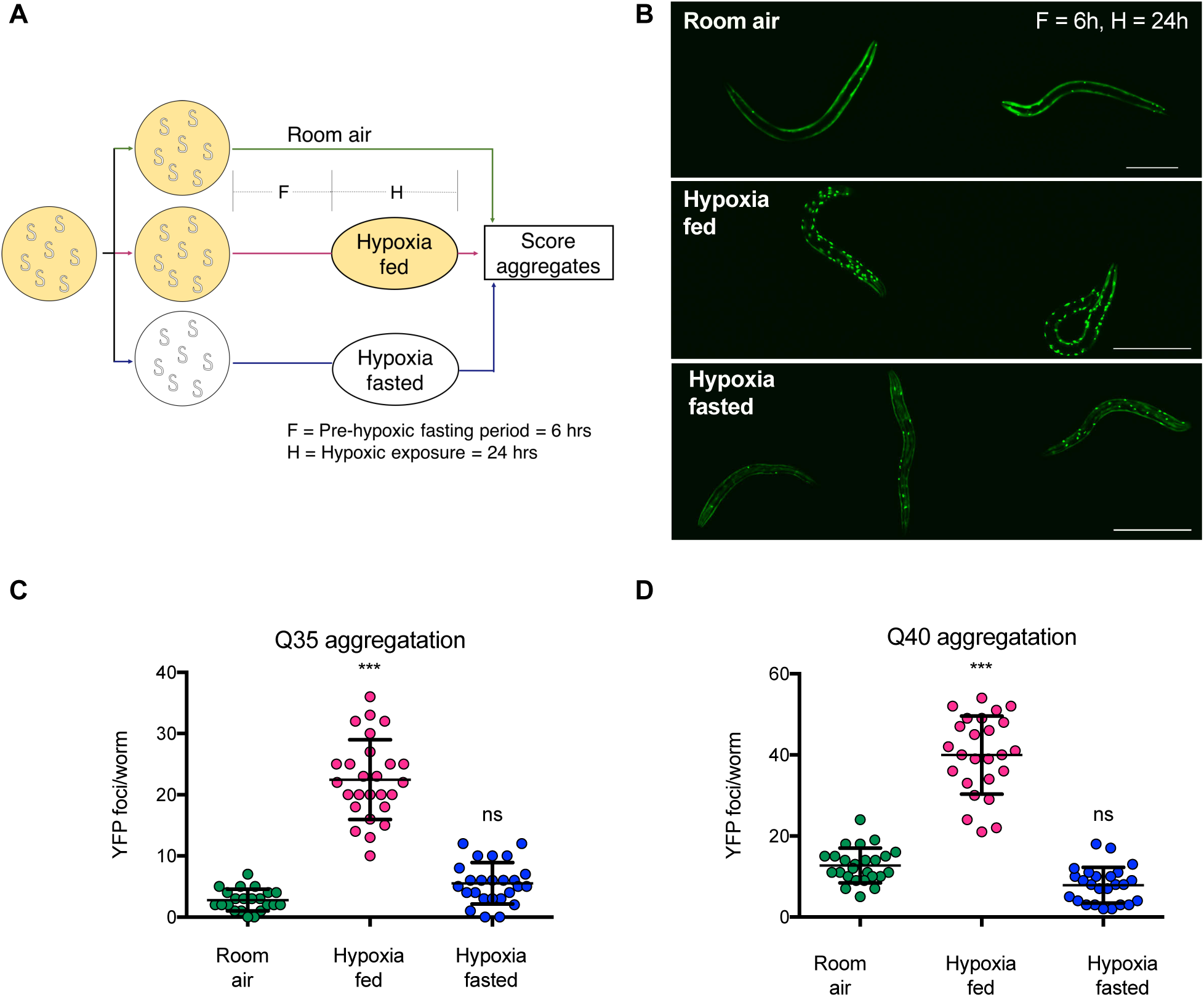
Fasting protects against hypoxia-induced protein aggregation. **A.** Experimental Schematic. Cohorts of age-synchronized animals were split into three groups: the first was maintained on food in room air, the second was maintained on food before and during exposure to hypoxia, and the third was removed from food before exposure to hypoxia. Fasting is indicated by white plates, yellow plates indicate animals on food. F= the duration of fasting (h) before hypoxia; H = duration of hypoxia (h). Unless otherwise noted, aggregates were counted immediately upon removal from hypoxia. **B.** Representative images of Q40∷YFP animals from cohorts of animals maintained in room air, exposed to hypoxia on food (hypoxia fed), or exposed to hypoxia while fasted (hypoxia fasted). F=6h, H=24h. Scale bars = 100μm**. C-D.** Aggregation measurements for L4 Q35∷YFP (**C**) and L1 Q40∷YFP (**D**) animals exposed to hypoxia on food (fed, magenta) or after removal from food (fasted, blue). Controls remained in room air (green). Data from one representative experiment is shown. Each experiment was repeated at least 3 times. Each circle is the number of YFP foci in a single animal, the mean is indicated by the line, and error bars are the standard deviation. Statistical comparisons were made between animals exposed to hypoxia and controls maintained in room air. Significance: *** *p* < 0.001; ns, not significant.

We originally chose to fast animals for 6h before exposure to hypoxia to allow animals time to alter gene expression [13], and this period of time off of food is sufficient to deplete stored glycogen as measured by iodine staining (DLM unpublished). However, there is no evidence to suggest that the protective effects of fasting in hypoxia requires changes in gene expression or glycogen stores. Therefore we next measured how long of a fasting period was required to mitigate the effects of hypoxia on aggregation of polyglutamine proteins.

To determine the pre-hypoxia fasting duration required to protect against HIPA, we removed Q35∷YFP animals from food for varying lengths of time before being exposed to hypoxia (as diagrammed in Fig. 2A). We found that animals removed from food immediately before exposure to hypoxia developed significantly fewer YFP foci in hypoxia as compared to controls that remained on food in hypoxia (Fig. 2A, 6h fed compared to fed). We conclude that extended fasting before exposure to hypoxia is not required to prevent HIPA. Instead, our data show that the protective effects of fasting occur very rapidly. In fact, the full protection against HIPA is realized with only 2h fasting before exposure to hypoxia (Fig. 2A). These results suggest that at least some of the protective effects of fasting are due to the absence of food directly, rather than metabolic changes or alterations in gene expression that occur during fasting prior to the hypoxic insult.

**Figure 2:**
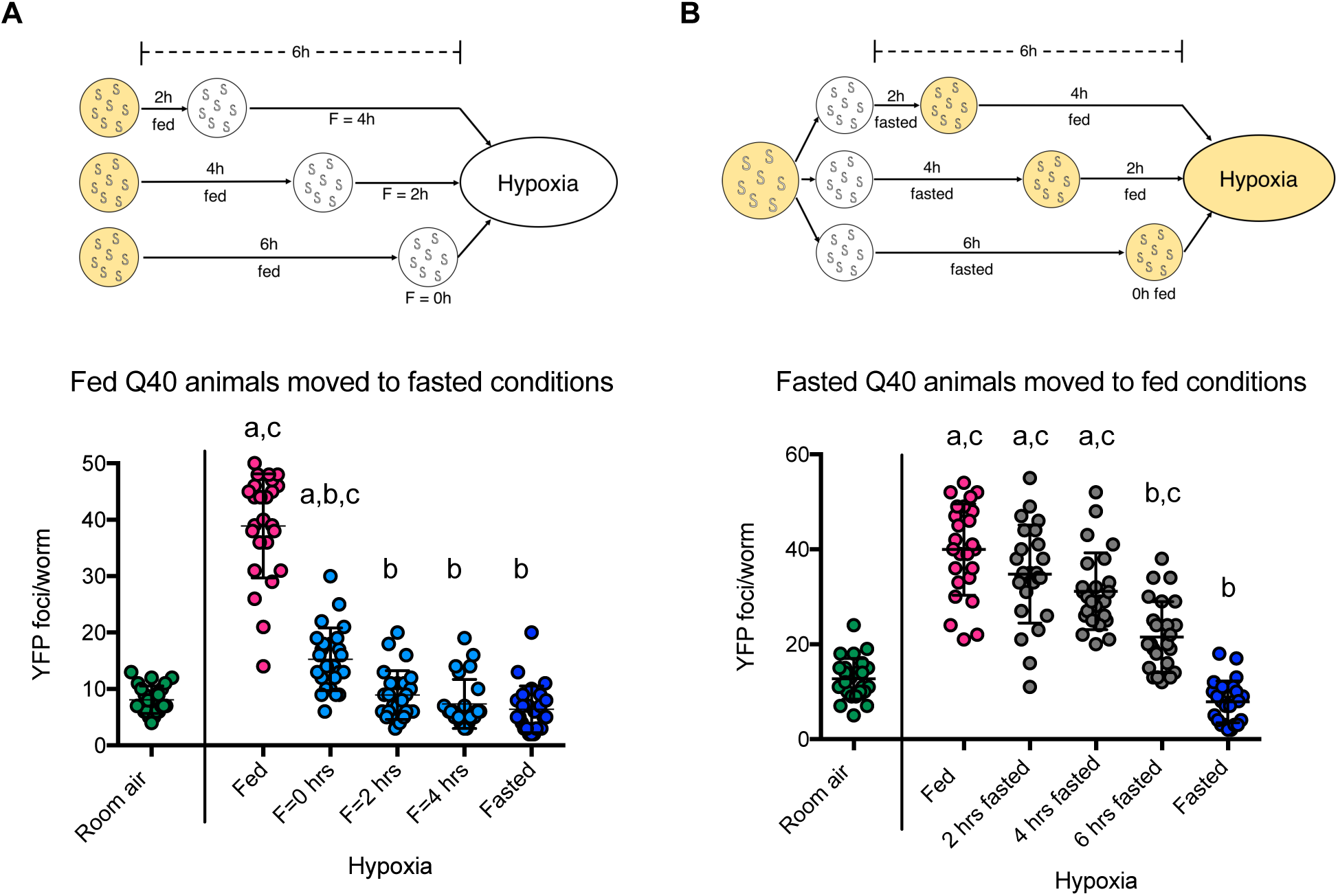
Fasting protection against HIPA is quickly induced and reversed. **A.** Effect of fasting occurs rapidly in hypoxic conditions. Cohorts of L1 Q40∷YFP animals were removed from food before exposure to hypoxia (F = 0, 2, or 4 h; H=24 h). All animals were off of food when exposed to hypoxia and the number of foci was scored immediately upon removal from hypoxia (cyan). Controls remained in room air (green), were continuously on food (fed, magenta), or were fasted for a full 6 h before hypoxia (fasted, blue). Data from one representative experiment is shown. Each experiment was repeated at least 3 times. Each circle is the number of YFP foci in a single animal, the mean is indicated by the line, error bars are the standard deviation. Significance was calculated using a Kruskal-Wallis test and Dunn’s multiple comparisons post hoc analysis. Significant differences (*p* < 0.05) in aggregation between conditions are indicated by letters above each group as follows: a - significantly different from room air controls; b - significantly different from fed hypoxic controls; c - significantly different from fasted hypoxic controls. **B.** Fasting before exposure to hypoxia improves proteostasis. As shown in the schematic above the graph, cohorts of L1 Q40∷YFP animals were removed from food 6h before exposure to hypoxia, and fasted for 2, 4, or 6 h before being returned to food. All cohorts were on food when exposed to hypoxia (H=24 h). The number of foci was scored immediately upon return to room air (gray). Controls remained in room air (green), were continuously on food and exposed to hypoxia (fed, magenta), or were not returned to food before hypoxia (fasted, blue). Data from one representative experiment is shown. Each experiment was repeated at least 3 times. Each circle is the number of YFP foci in a single animal, the mean is indicated by the line, error bars are the standard deviation. Statistical comparisons were made between animals fasted for the indicated amount of time and controls maintained in room air, fed controls exposed to hypoxia after being continuously on food, and fasted controls that were not returned to food before hypoxia. Significance was calculated using a Kruskal-Wallis test and Dunn’s multiple comparisons post hoc analysis. Significant differences (*p* < 0.05) in aggregation between conditions are indicated by letters above each group as follows: a - significantly different from room air controls; b - significantly different from fed hypoxic controls; c - significantly different from fasted hypoxic controls.

Work in other systems has shown that fasting can have a protective effect that persists even after animals are returned to food (Robertson and Mitchell 2014). To further explore the requirements for fasting to protect against HIPA we next asked whether the protective effects of fasting against HIPA could be reversed. In these experiments (Fig. 2B), we began fasting animals 6h before exposure to hypoxia but then returned the animals to food prior to initiation of hypoxia. We observed that animals fasted for a full 6h and then returned to food immediately before exposure to hypoxia (Fig. 2B, 6h fasted) developed significantly more YFP foci than animals that were fasted for 6h and then exposed to hypoxia in the absence of food (Fig 2B, fasted), suggesting that the nutritional context of an animal as it experiences hypoxia is able to mediate the effect of hypoxia on proteostasis. Furthermore, we found no protection from HIPA if animals were fasted for 4h, but then fed for 2 h before exposure to hypoxia (Fig. 2B, 4h fasted), even though 4h of fasting was sufficient for complete protection against HIPA in the absence of food (Fig. 2A, 2h fed). This result indicates that the protective effects of fasting are fully reversed within 2h of return to food. We conclude that the protective effects of fasting in hypoxia are rapidly reversed.

**Figure 3:**
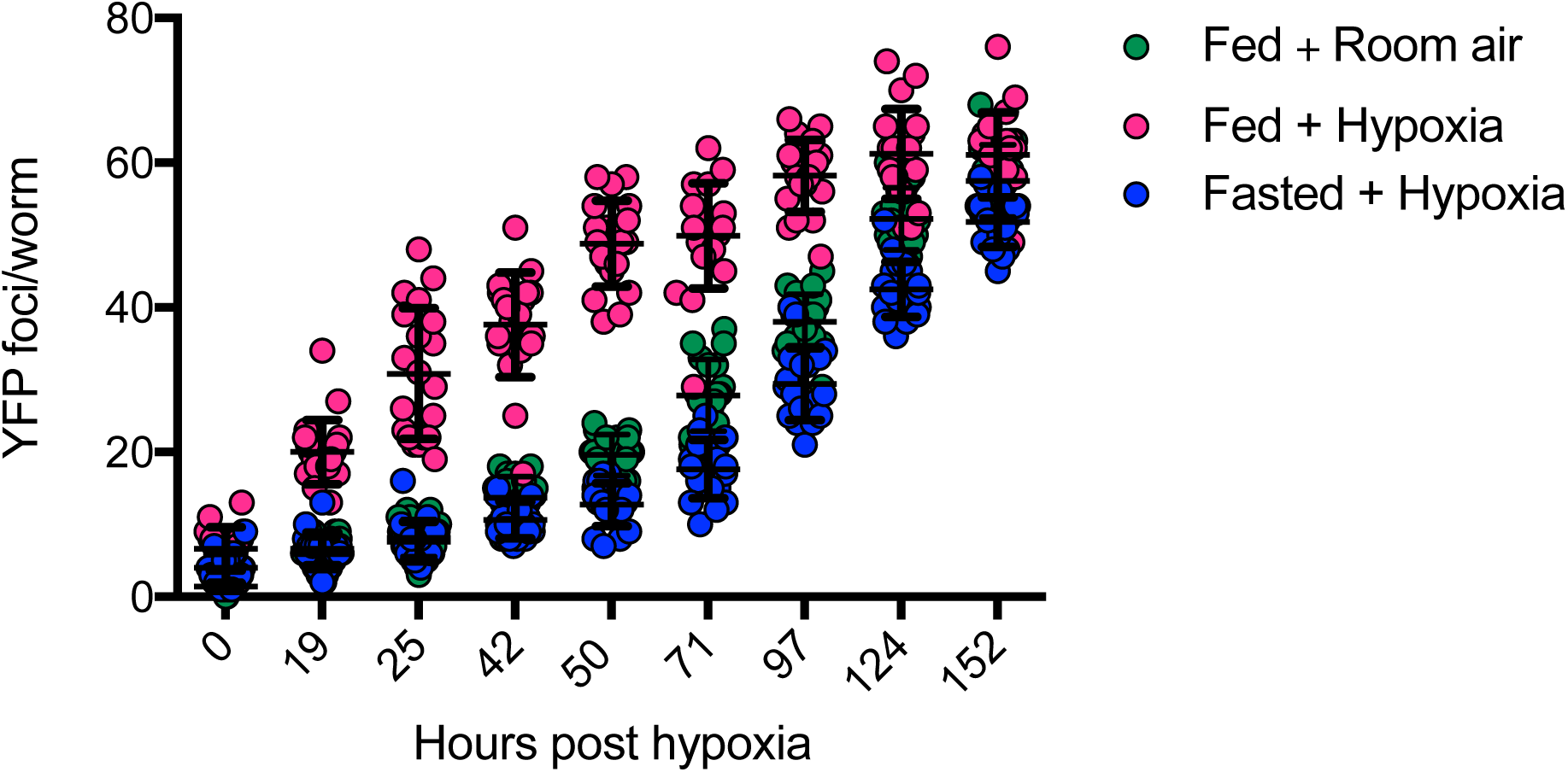
Fasting protects against long-term effects of hypoxia on proteostasis. Cohorts of L4 Q35∷YFP animals were exposed to hypoxia (H=10 h) on food (magenta) or fasted (blue, F=6h). Controls remained in room air on food (green). The number of YFP foci was scored after return to room air as indicated. Data from one representative experiment is shown. The experiment was repeated at least 3 times. Each cohort included at least 20 animals per time point.

**Figure 4.**
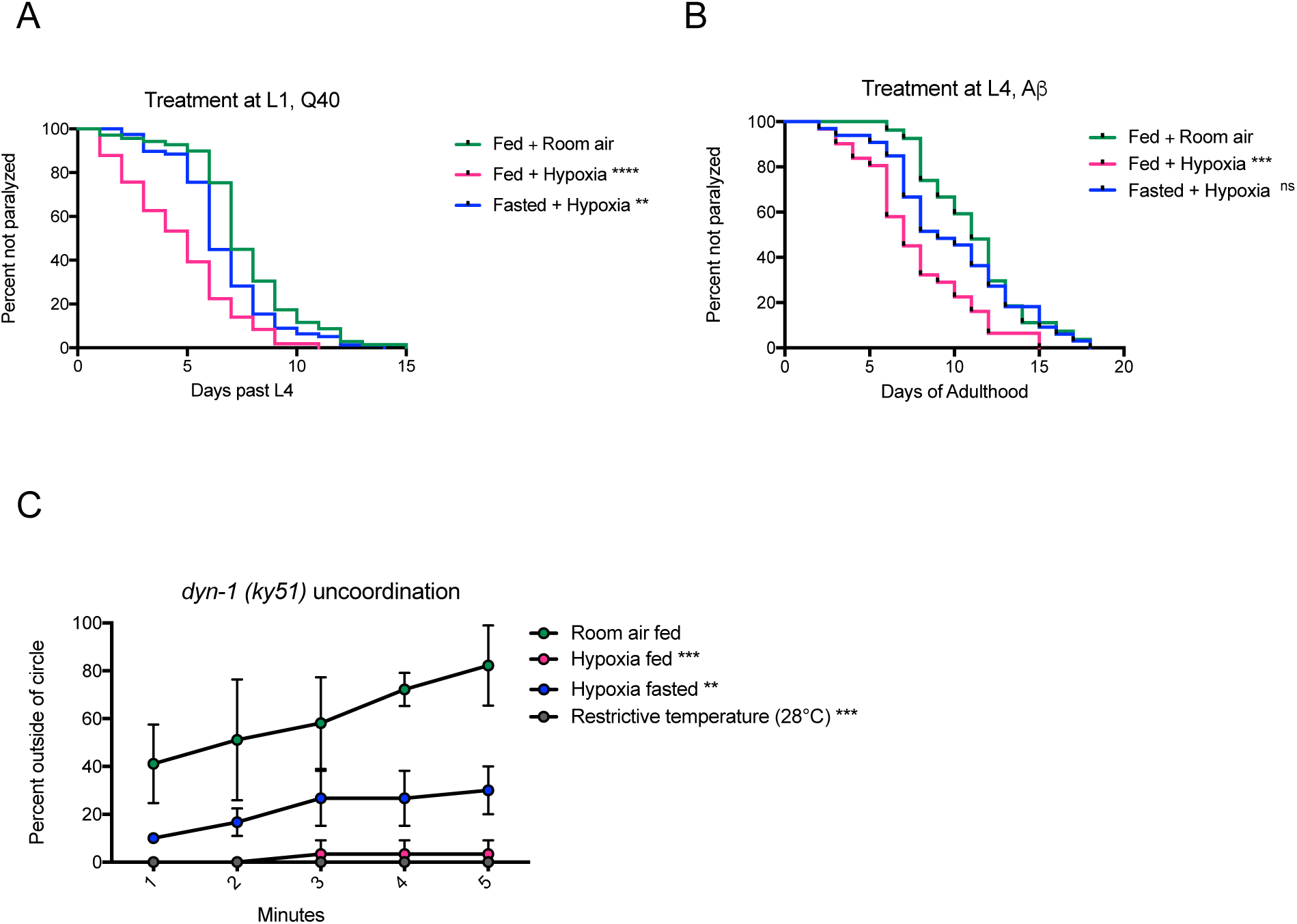
Fasting has general protective effects against hypoxia-induced defects in proteostasis. **A.** Fasting protects against toxicity of Q40∷YFP. Cohorts of L1 animals expressing Q40∷YFP were exposed to hypoxia on food (magenta), or fasted (blue) before exposure to hypoxia (F=6h, H=24 h). Paralysis was scored after return to room air, beginning the first day of adulthood. Controls remained on food in room air (green). Data from one representative experiment is shown, each cohort included at least 70 animals. Each experiment was repeated at least 3 times. Significance was calculated using a Log-rank (Mantel-Cox) test with a Bonferroni correction for multiple comparisons. Statistical comparisons were made between animals exposed to hypoxia and animals maintained in room air. **** *p* < 0.0001; ** *p* < 0.01. **B**. Fasting protects against toxicity of Aβ_1-42_. Cohorts of L4 animals expressing Aβ_1-42_ were exposed to hypoxia on food (magenta) or fasted (blue) before exposure to hypoxia (F=6h, H=24h). Paralysis was scored after return to room air, beginning at the first day of adulthood. Controls remained on food in room air (green). Data from one representative experiment is shown, each cohort included at least 70 animals. Each experiment was repeated at least 3 times. Significance was calculated using a Log-rank (Mantel-Cox) test with a Bonferroni correction for multiple comparisons. Statistical comparisons were made between animals exposed to hypoxia and animals maintained in room air. *** *p* < 0.001. **C.** Fasting protects against hypoxia effects on metastable DYN-1. Temperature-sensitive *dyn-1(ky51)* mutant animals were exposed to hypoxia at the permissive temperature on food (magenta), or after fasting (blue). Controls remained on food in room air at the permissive temperature (green) or on food at the non-permissive temperature (28°C, gray). Paralysis was scored 1h after return to room air. Average data from 3 independent experiments is shown, each cohort included 10 animals. Significance was calculated using a repeated measures two-way ANOVA and Dunnett’s multiple comparisons test. Statistical comparisons were made between animals exposed to hypoxia or animals maintained at the restricted temperature and animals maintained in room air. Significance: *** *p* < 0.001; ** *p* < 0.01

Shorter exposures to hypoxia that do not immediately increase the number of polyglutamine protein aggregates still disrupt long-term proteostasis, as evidenced by the increased rate of age-associated protein aggregation after return to room air [10]. We therefore asked whether fasting could protect against these long-term proteostasis deficits in addition to HIPA. We exposed Q35∷YFP L4 animals to hypoxia for only 10h either in the fed state or after fasting for 6h (F = 6 hours, H = 10 hours as per Fig. 1A). Control animals remained on food in room air. Immediately after this short hypoxic exposure, there was no observed increase in the number of YFP foci in animals exposed to hypoxia regardless of whether food was present (Fig. 3, 0 hours post-hypoxia). As expected, the animals exposed to hypoxia in the fed state accumulate aggregates faster than control animals. In contrast, animals exposed to hypoxia while fasted accumulate YFP foci at the same rate as control animals. These data indicate that fasting both prevents HIPA and protects against the long-term effects on proteostasis induced by a short exposure to hypoxia.

The cellular role of protein aggregates is controversial, with some reports finding a protective role and others suggesting a cytotoxic effect [14]. We have previously shown that aggregates induced by hypoxia are cytotoxic, resulting in accelerated paralysis after animals are returned to room air [10]. We therefore next asked if fasting would protect against increased proteotoxicity in addition to HIPA. To address this, we exposed cohorts of L1 Q40∷YFP animals to hypoxia for 24 hours while fed or fasted, then returned the animals to room air and measured the onset of paralysis in each cohort. We found that fasting slowed the rate at which paralysis developed relative to animals exposed to hypoxia while fed (Fig. 4A). This result indicates that fasting protects against hypoxic effects of increased protein aggregation and proteotoxicity.

We next sought to determine whether fasting’s protective effects on proteostasis extend to other models of proteotoxicity. Human amyloid β (Aβ)_1-42_ peptide expressed in the body wall muscles of *C. elegans* results in cytoplasmic plaque formation, with a subsequent phenotype of progressive paralysis [15]. *C. elegans* expressing Aβ_1-42_ in their body wall muscles become paralyzed more quickly when they are exposed to hypoxia [10]. We found that this effect of hypoxia was reversed by fasting, as the rate that paralysis develops is slowed if animals expressing Aβ_1-42_ are exposed to hypoxia while fasting (Fig. 4B). Because Aβ_1-42_ and Q40∷YFP are both expressed in body wall muscles, we also evaluated if fasting protected animals expressing a metastable version of the neuronal dynamin protein DYN-1 from the effects of hypoxia. The *dyn-1(ky51)* mutant contains a temperature-sensitive (ts) mutation, such that the DYN-1 protein is functional and *dyn-1(ky51)* mutant animals exhibit wild-type motility at the permissive temperature (20°C), but become uncoordinated at the restrictive temperature (28°C) due to improper folding of the DYN-1 protein [16]. Genetic and environmental factors that disrupt proteostasis, including hypoxia, prevent the proper folding of the DYN-1 protein at the permissive temperature, thereby rendering the *dyn-1(ky51)* animals uncoordinated [17, 10]. Similar to our experiments with Q40∷YFP and Aβ_1-42_, we found that fasting *dyn-1(ky51)* mutant animals before exposure to hypoxia results in a partial rescue of hypoxia-induced uncoordination at the permissive temperature (Fig. 4C). Together, our results suggest that fasting has a general protective effect against proteostasis defects induced by hypoxia, and that this protective effect is not specific to a particular tissue, developmental stage, or misfolded/aggregation prone model.

Dysregulation of insulin-like signaling (IIS) has been tied to protein aggregation and neurodegeneration in a number of model organisms [18]. As the IIS pathway links food availability to growth, development, stress resistance, and aging, we hypothesized that changes in IIS could explain how fasting modulates the effect of hypoxia on proteostasis. The IIS pathway is widely conserved in metazoans [19]. We therefore explored the hypothesis that IIS would mediate the effects of fasting to prevent HIPA.

We first looked at the localization of DAF-16∷GFP in animals exposed to hypoxia to determine if IIS is active in hypoxia. DAF-16 is the *C. elegans* orthologue of the FOXO transcription factor. When active, the insulin/IGF-like receptor DAF-2 initiates a phosphorylation cascade that results in the phosphorylation and nuclear exclusion of DAF-16 protein [20, 21]. Conversely, when nutrients are scarce, DAF-16 remains unphosphorylated by upstream kinases and is able to enter the nucleus and bind to its target genes [20, 22]. We found that DAF-16∷GFP remained diffuse and cytoplasmic in control worms maintained in room air on food (Fig 5B, 5C), but accumulated in the nucleus of animals that were removed from food in room air (Fig. 5B, 5C) or were exposed to hypoxia on food (Fig. 5B, 5C). These results suggest that IIS activity is reduced by fasting and hypoxia, consistent with previous reports [23, 24]. Surprisingly, DAF-16∷GFP did not accumulate in the nuclei of animals exposed to hypoxia after fasting (Fig 5B, 5C), despite hypoxia and fasting both individually resulting in nuclear accumulation.

**Figure 5.**
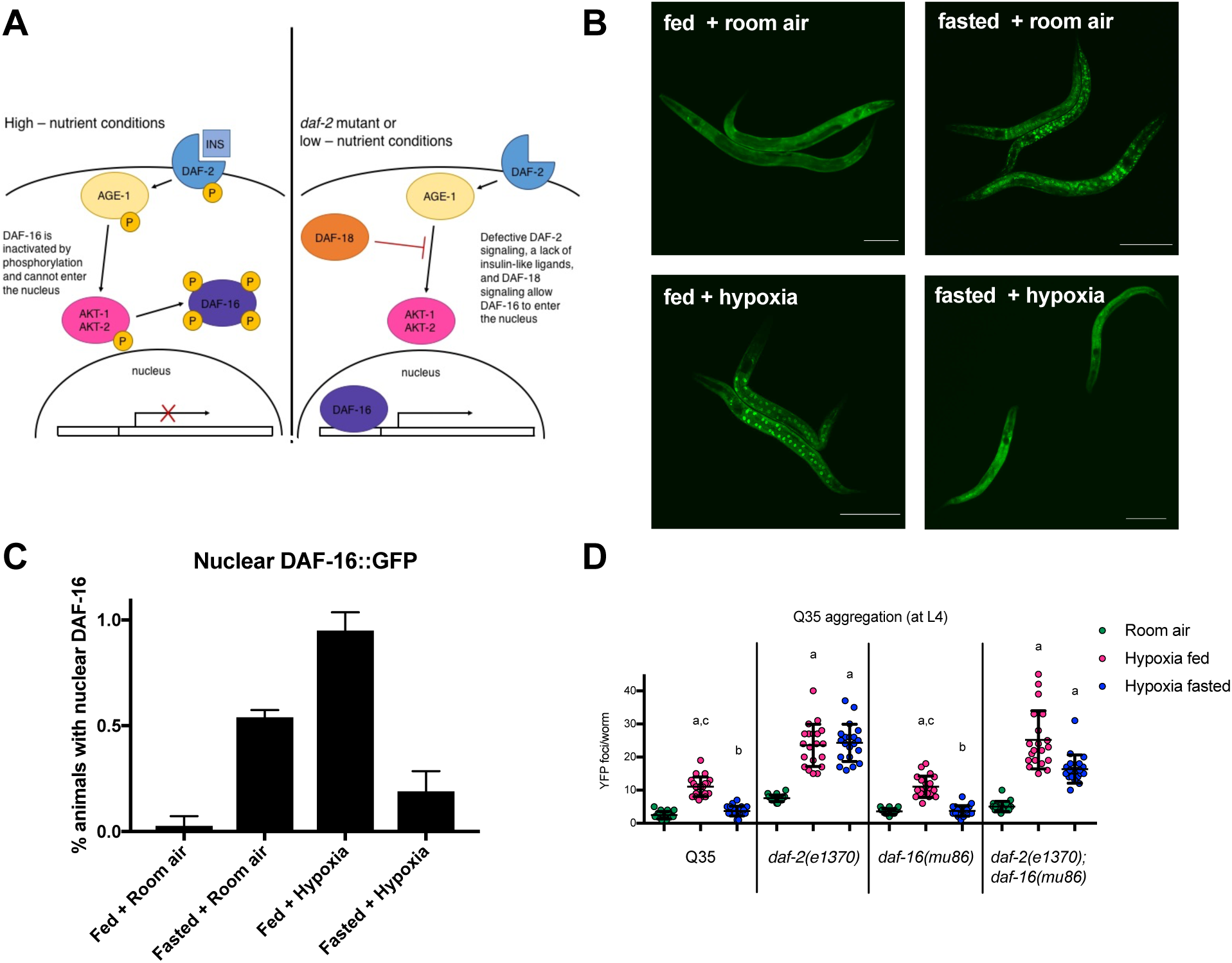
The insulin-like signaling pathway is required for fasting protection. **A** Schematic of key insulin-signaling pathway members in *C. elegans*. Under nutrient-rich conditions, insulin-like peptides bind to the insulin receptor DAF-2, initiating a phosphorylation cascades that ultimately leads to the phosphorylation of the FoxO transcription factor DAF-16, excluding it from the nucleus. Conversely, when nutrients are scarce, DAF-16 remains unphosphorylated and is able to enter the nucleus and bind to its target genes. **B.** DAF-16 is not localized to the nucleus in fasted animals exposed to hypoxia. Cohorts of 20 DAF-16∷GFP animals were maintained in room air on food for 24 hrs (fed + room air), fasted in room air for 24 hrs (fasted + room air), exposed to hypoxia for 24 hrs on food (fed + hypoxia), or exposed to hypoxia after fasting (fasted + hypoxia; F=6h, H=24hr). Scale bars = 100μm. **C** Quantification of DAF-16∷GFP nuclear accumulation. The percent of animals with nuclear GFP was scored immediately post hypoxia. Average data from 3 independent experiments is shown. The bar height indicates the mean. Error bars are the standard deviation. **D** Fasting does not protect *daf-2* mutants against HIPA. Aggregation measurements (F=6h, H=24h) for L4 Q35∷YFP animals with mutations in *daf-2(e1370), daf-16(mu86)*, and the *daf-2(e1370); daf-16(mu86)* double mutant. Animals were maintained on food in room air (room air, green), were exposed to hypoxia on food (fed, magenta), or were exposed to hypoxia after removal from food (hypoxia fasted, blue). Each circle is the number of YFP foci in a single animal, the mean is indicated by the line, error bars are the standard deviation. Data from one representative experiment is shown. Each cohort included at least 20 animals, and each experiment was repeated at least 3 times. Significance was calculated using a Kruskal-Wallis test and Dunn’s multiple comparisons post hoc analysis. Significant differences (*p* < 0.05) in aggregation for a given strain between conditions are indicated by letters above each group as follows: a - significantly different from room air controls; b - significantly different from fed hypoxic controls; c - significantly different from fasted hypoxic controls.

These DAF-16∷GFP localization patterns led us to interrogate requirements for DAF-16 and the upstream IIS receptor DAF-2 in mediating fasted and fed responses to hypoxia. To this end, we crossed the Q35∷YFP transgene into *daf-2(e1370)* and *daf-16(mu86)* backgrounds. The fact that DAF-16∷GFP is localized to the nucleus in fed animals exposed to hypoxia suggests the possibility that DAF-16 facilitates HIPA. We found that *Q35∷YFP; daf-16(mu86)* mutant animals exhibit robust HIPA on food (Fig 5D), indicating that DAF-16 is not required for HIPA despite its nuclear accumulation in fed hypoxic animals. We also asked if there was a genetic requirement for the IIS receptor DAF-2. Our data indicate that IIS does not mediate the effects of hypoxia on proteostasis in fed animals, as *Q35∷YFP; daf-2(e1370)* mutant animals exhibit robust HIPA when fed (Fig. 5D). Thus, neither DAF-16 nor DAF-2 activities are required for HIPA in fed animals.

Given the IIS-independent nature of HIPA in fed animals, we next investigated whether fasting protection requires IIS. We discovered that DAF-2, but not DAF-16 is required for fasting protection against HIPA. Fasting protects the *Q35∷YFP; daf-16(mu86)* similar to wild-type (Fig 5D); however, we observe significant HIPA when *Q35; daf-2(e1370)* and *Q35; daf-2(e1368)* mutant animals are exposed to hypoxia when fasted (Fig 5D and Supplemental Fig. 1). These results show that protective effects of fasting in hypoxia require DAF-2, but not DAF-16. This is consistent with our observation that DAF-16∷GFP is not localized to the nucleus in fasted animals exposed to hypoxia (Fig 5B, 5C).

We found that the insulin/IGF-like receptor DAF-2 mediates the protective effects of fasting on HIPA, while the the FOXO transcription factor DAF-16 is not required for protection. Given this finding, we also checked the DAF-16∷GFP localization pattern in worms with a *daf-2(e1370)* mutation. These mutants have constitutively nuclear DAF-16 in the fed state due to decreased signaling through the IIS pathway [20]. Since DAF-16∷GFP is not localized to the nucleus in fasting-protected wild-type animals exposed to hypoxia, we sought to investigate whether the nuclear localization of DAF-16 in *daf-2(e1370)* mutants, which are not protected by fasting, would be altered by hypoxia. We found that DAF-16∷GFP is fully nuclear in all conditions, including fasted hypoxia, in these animals (Supplemental Fig. 2).

In *C. elegans*, DAF-16 mediates the effects of decreased signaling through DAF-2. Mutations in *daf-16* suppress most *daf-2* mutant phenotypes including increased lifespan, enhanced dauer formation, increased fat storage, reproductive delays, and increased resistance to heat and oxidative stress. [25, 26]. This coupled with the nuclear localization of DAF-16∷GFP in *daf-2* mutants led us to hypothesize that *daf-16* would be required for the HIPA in fasted *Q35; daf-2(e1370)* mutant animals. While *Q35; daf-16(mu86)* mutant animals were protected from HIPA by fasting similar to wild-type controls, *Q35; daf-2(e1370);daf-16(mu86)* animals still exhibit significant HIPA when fasted (Fig. 5D). These results indicate that DAF-2 mediates the effects of fasting to prevent HIPA at least partly independently of DAF-16.

## Discussion

This study illustrates the power of fasting to ameliorate the deleterious effects of hypoxia on proteostasis. These findings are consistent with phenomena that have been observed in mammals – fasting mice for a single day increases survival after kidney ischemia and also reduces ischemic damage to the liver [27]. Our results suggest that the nutritional milieu present at the onset of hypoxia can dictate the effect of hypoxia on proteostasis, as fasting protection against hypoxia can be induced quite quickly. Animals that are removed from food immediately before hypoxia are protected against HIPA to a significant degree, even after being maintained on food for the entire pre-hypoxic period. This implies that worms are integrating information about their environment, including nutrient availability, right as they sense hypoxia. The importance of the nutritional environment of the animal as it experiences hypoxia is further supported by the fact that we also see a rapid reversal of fasting protection. Worms fasted for six hours but that are moved onto food immediately preceding hypoxia are not as protected against HIPA compared to worms that were fasted and remained off of food for the duration of hypoxia. The speed with which fasting protection can be induced and reversed indicates that protection cannot be explained solely by changes in gene expression resulting in a hypoxia-resistant pre-adapted state. Furthermore, the rapidity with which fasting protection can be reversed suggests that altered gene expression or metabolism resulting from the fasting period is alone insufficient to protect against HIPA. Although *C. elegans* enter a reproductive and developmental diapause in 0.1% oxygen [28], the protection conferred by fasting does not represent a simple delay in the onset of proteostasis decline due to the time spent in hypoxia. Rather, fasting provides long-term protection against the accrual of protein aggregates and toxicity even after the return to room air.

We found that IIS mediates fasting protection against HIPA. Notably, IIS is not required for the fed response to hypoxia, as fed IIS mutants show increased aggregate levels comparable to wild-type animals. In worms and flies, mutations in the insulin receptor are generally considered protective against hypoxia. In *C. elegans, daf-2* mutants have a hypoxia-resistant phenotype, displaying reduced muscle and neuronal cell death following hypoxia [29, 30], while flies with defective insulin signaling (mutations in the insulin receptor *InR*, or *Chico*, the insulin receptor substrate) are protected against anoxia/reoxygenation injury [31]. The *daf-2* phenotype uncovered here is therefore distinct in that these mutants are sensitive to hypoxia in the fasted state, with fasted *daf-2* mutant animals exhibiting increased HIPA compared to wild-type controls. These results contradict the *a priori* expectation that *daf-2* mutants might be resistant to hypoxia even in the fed state due to their inability to detect insulin-like peptides.

Mammalian systems offer precedents of insulin receptor mutations causing sensitivity to hypoxic stress. Knockdown of neuronal insulin-like growth factor 1 receptor (IGF-1R) exacerbates hypoxic injury and increases mortality in mice [32], and IGF-1R is required in order for IGF-1 to protect myocardial cell exposed to ischemia [33]. However, data on the role of mammalian IIS in response to hypoxia are mixed, and are complicated by the fact that different types of insulin receptors mediate distinct cellular functions [34]. As such, the simplified *C. elegans* IIS system may be useful for understanding contextual inputs that alter IIS outputs.

DAF-16 is believed to be the main nexus of IIS [20, 35-37], which makes the DAF-2-dependent, but DAF-16-independent nature of the protective effect of fasting described here unusual in *C. elegans*. Decreased DAF-2 activity results in phenotypes such as increased lifespan, reproductive delays, and increased resistance to heat and oxidative stress, all of which require DAF-16 [26]. However, a few other examples exist in the literature of DAF-2 dependent, DAF-16 independent phenomena: dauer formation at 27°, meiotic progression of oocytes, salt chemotaxis learning, and regulation of the dao-3 and hsp-90 genes [38-42]. In chemotaxis learning, DAf-2 acts on learning through phosphatidylinositol 3,4,5-triphosphate (PIP_3_), but not DAF-16. Similar to these studies, fasting-mediated protection against HIPA supports the existence of downstream targets of DAF-2 separate from DAF-16 that are capable of influencing stress responses and proteostasis.

## Materials and methods

### C. elegans strains and methods

Animals were maintained on nematode growth media (NGM) with OP50 *E. coli* at 20°C (Brenner, 1974). See Supplementary Table S6 for worm strains. Strains were obtained from the *Caenhorabditis* Genetics Center at the University of Minnesota. Double and triple mutants were generated using standard genetic techniques, and genotypes were verified using PCR.

### Construction of hypoxic chambers

Hypoxic conditions were maintained using continuous flow chambers, as described in Fawcett et al. 2012. Compressed gas tanks (1000 ppm O_2_ balanced with N_2_) were Certified Standard (within 2% of target concentration) from Airgas (Seattle, WA). Oxygen flow was regulating using Aalborg rotameters (Aalborg Intruments and Controls, Inc., Orangeburg, NY, USA). Hypoxic chambers (and room air controls?) were maintained in a 20°C incubator for the duration of the experiments.

### YFP∷polyQ aggregation assays

Synchronous cohorts of L1 YFP∷polyQ_40_ animals were generated by either bleaching first-day adult animals in a 20% alkaline hypochlorite solution or allowing first-day adult animals to lay eggs for 1-2 hrs on seeded NGM plates. The adults were then removed, and the plates were incubated at 20°C. The next morning, cohorts of hatched L1 larvae were suspended in M9 and mouth-pipetted to new NGM plates for hypoxic exposure. Synchronous cohorts of L4 YFP∷polyQ_35_ animals were generated by picking L4 animals from well-fed, logarithmically growing populations.

Cohorts of 25-35 YFP∷polyQ animals were exposed to hypoxia for approximately 24 h at 20°C on unseeded 3 cm NGM plates with 40mg/mL carbenicillin or NGM plates seeded with live OP50 food. Plates were ringed with palmitic acid (10mg/mL in ethanol), creating a physical barrier around the edge of each plate to discourage animals from leaving the surface of the agar.

To quantify the number of YFP foci, worms were mounted a 2% agar pad in a drop of 50mM sodium azide as anesthetic. Control experiments showed that azide did not affect the aggregation of YFP∷polyQ_35_ or YFP∷polyQ4_0_ (Moronetti Mazzeo et al. 2012). YFP foci were identified and quantified as described in Morley et al. (2002) and Silva et al. (2011). A Nikon 90i fluorescence microscope with the YFP filter and 10x objective (Nikon Instruments Inc., Melville, NY, USA) was used to visualize and quantify aggregates. In all experiments, the number of aggregates was counted blind to treatment and genotype. Statistical significance was evaluated by calculating P-values between conditions using a Kruskal-Wallis test and Dunn’s multiple comparisons post hoc analysis in GraphPad Prism version 7.0c for Mac OSX (GraphPad Softare, San Diego, California, USA) In all cases, P < 0.05 was considered statistically significant.

### Paralysis and uncoordination assays of proteotoxicity

Animals expressing Aβ_1-42_ or YFP∷polyQ_40_ were exposed to 1000 ppm O_2_ for 24 at 20°C as L4 or L1, respectively. For both, animals were grown on seeded NGM plates until 6 hrs before hypoxic exposure, at which point fasted animals were transferred to unseeded NGM plates, where they remained until the end of the hypoxic exposure. Fed animals were transferred to new seeded NGM plates. After hypoxic exposure, all animals were returned to food and normoxia, and incubated at 20°C. Paralysis was scored daily. Worms were considered paralyzed if they failed to respond, other than with movement of the nose or pharyngeal pumping, when tapped with a platinum wire pick 3 consecutive times. Dead or bagged worms were censored from the experiment on the day of death/bagging. Paralyzed worms were removed from the plate on the day of paralysis. Live worms that were not paralyzed were moved to a new plate each day until all worms were scored as either paralyzed or dead. Statistical significance was calculated using Kaplan-Meier log-rank (Mantel-Cox) tests and a Bonferroni correction for multiple comparisons using GraphPad Prism version 7.0c for Mac OSX (GraphPad Softare, San Diego, California, USA).

### DAF-16∷GFP localization

Synchronous cohorts of L2 animals expressing DAF-16∷GFP were exposed to hypoxia for 24 h at 20°C on unseeded unseeded 3 cm NGM plates with 40mg/mL carbenicillin or NGM plates seeded with live OP50 food. Plates were ringed with palmitic acid (10mg/mL in ethanol), creating a physical barrier around the edge of each plate to discourage animals from leaving the surface of the agar. To visualize the localization of DAF-16∷GFP, worms were mounted a 2% agar pad in a drop of 10mM levamisole as anesthetic. A Nikon 90i fluorescence microscope with the GFP filter and 10x objective (Nikon Instruments Inc., Melville, NY, USA) was used to visualize DAF-16∷GFP. For quantification, percent of animals with nuclear GFP was scored immediately after removal from hypoxia. In all experiments, the GFP localization was scored blind to treatment and genotype. Statistical significance was evaluated by calculating P-values between conditions using a Kruskal-Wallis test and Dunn’s multiple comparisons post hoc analysis in GraphPad Prism version 7.0c for Mac OSX (GraphPad Softare, San Diego, California, USA). P < 0.05 was considered statistically significant.

## Acknowledgements

We

