## Supplementary Materials for "Fasting prevents hypoxia-induced defects of proteostasis in *C. elegans*"

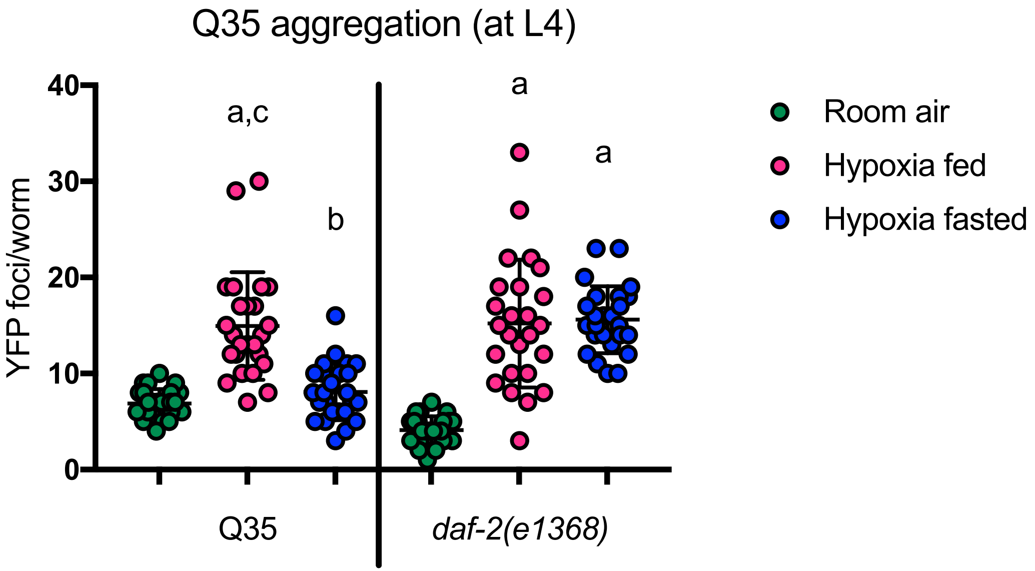


Supplemental Figure 1. Fasting dooes not protect *daf-2(e1368)* mutants against HIPA. Aggregation measurements (F=6h, H=24h) for L4 *daf-2(e1368)* Q35::YFP animals. Animals were maintained on food in room air (green), were exposed to hypoxia on food (magenta), or were exposed to hypoxia after removal of food (blue). Each circle is the number of YFP foci in a single animal. The mean in indicated by the line, error bars are the standard deviation. Data from one representative experiment is shown. Each cohort included at least 20 animals, and the experiment was repeated at least 3 times. Significance was calculated using a Kruskal-Wallis test and Dunn's multiple comparisons post hoc analysis. Signficant differences ($p$<0.05) in aggregation for a given strain between conditions are indicated by letters above each group as follows: a - significantly different from room air controls; b - significantly different from fed hypoxic controls; c - significantly different from fasted controls.
