## Supplementary Materials for "Fasting prevents hypoxia-induced defects of proteostasis in *C. elegans*"

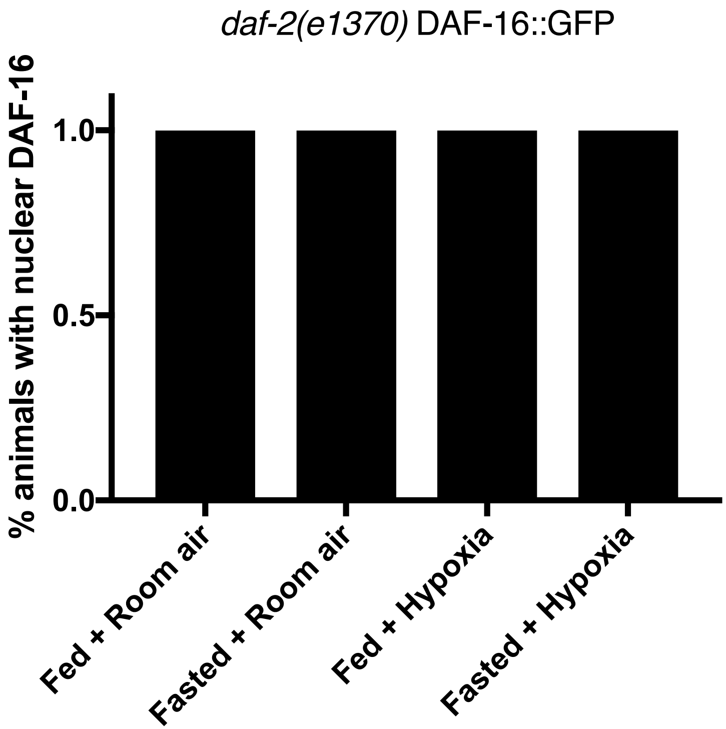


Supplemental Figure 2. DAF-16::GFP in *daf-2(e1370)* mutants is localized to the nucleus in all fasted, fed, and hypoxic conditions. Cohorts of 20 *daf-2(e1370)* mutants expresseing DAF-16::GFP were maintained in room air on food for 24 hours (Fed + Room air), fasted in room air for 24 hours (Fasted + Room air), exposed to hypoxia for 24 hours on food (Fed + Hypoxia), or exposed to hypoxia after fasting (Fasted + Hypoxia; F=6h; H=24H). The percent of animals with nuclear GFP was scored immediately post hypoxia. Average data from 3 independent experiments is shown. The bar height indicates the mean. Error bars (present, but not visible) are the standard deviation.
