## Supplementary Materials for "Fasting prevents hypoxia-induced defects of proteostasis in *C. elegans*"

| Strain | Genotype | Reference |
| --- | --- | --- |
| AM140 | rmls132 [unc-54p::Q35::YFP] | Saytal et al. 2000 |
| AM141 | rmls133 [unc-54p::Q40::YFP] | Saytal et al. 2000 |
| CX51 | dyn-1 (ky51) | Clark et al. 1997 |
| CL2006 | dvls2 [pCL12(unc54/human Abeta peptide 1-42 minigene) + pRF4] | Link, 1995 |
| CB1370 | daf-2(e1370) | Kimura et al. 1997 |
| CF1038 | daf-16(mu86) | Lin et al. 1997 |
| TJ356 | zls356 [daf-16p::daf-16a/b::GFP + rol-6(su1006)] | Lin et al. 2001 |
| GR1895 | daf-2(e1370); mgls67 [daf-16p::daf-16::GFP + rol6(su1006)] | Riedel et al. 2013 |
|  | daf-2(e1370); Q35::YFP | * |
|  | daf-2(e1368); Q35::YFP |  |
|  | daf-16(mu86); Q35::YFP | * |
|  | daf-2(e1370); daf-16(mu86); YFP::Q35 | * |

* Strain was created by crossing AM140 with indicated genetic background. Mutant alleles were verified using PCR genotyping.
